# Bacterial proteome foundation model enhances functional prediction from enzymes to ecological interactions

**DOI:** 10.64898/2026.03.07.710335

**Authors:** Palash Sethi, Lucas A. Pereira, Juannan Zhou

## Abstract

Bacteria play fundamental roles in ecosystems, human health, and biotechnology. Although bacterial genome sequencing data have accumulated rapidly over the past decade, the metabolic and ecological functions carried out by most sequenced bacteria remain poorly understood, apart from a few well-studied taxa and traits. Establishing a general framework that comprehensively captures the relationship between bacterial genomes and the diverse biological functions they encode remains a major challenge, as it requires embedding individual genes within their broader genomic context and modeling their combined effects across complex biological pathways and networks. The difficulty is further compounded by the limited functional annotations available for most bacterial genomes. Here, we introduce BacPT, a proteome foundation model trained on tens of thousands of complete genomes spanning diverse bacterial taxa. BacPT captures both local and genome-wide information, enabling the generation of contextualized gene embeddings and functionally rich representations of the whole genome. We demonstrate the utility of BacPT across diverse prediction tasks spanning multiple biological scales. BacPT embeddings improve the prediction of enzyme activities, biosynthetic gene clusters, metabolic traits, and ecological interaction outcomes. Our results highlight that unsupervised deep learning applied at the scale of entire proteomes provides a powerful approach for characterizing gene interactions and mapping functional landscapes for bacteria.

## Introduction

The remarkable metabolic diversity of bacteria stems from the vast repertoire of enzymes, gene pathways, and regulatory networks shaped by billions of years of evolution [1–3]. They enable bacteria to thrive across a broad spectrum of environments and carry out critical ecological functions, such as nutrient cycling, symbiotic interactions, and pathogenesis [3–10]. Consequently, mapping the functional landscape encoded within bacterial genomes and linking genomic sequences to specific metabolic and ecological traits is fundamental to advancing disease biology, human health, synthetic biology, and microbial ecology [11, 12].

In recent years, advances in high-throughput sequencing technologies have led to a rapid expansion of bacterial genomic data, opening many exciting opportunities to explore the relationship between bacterial genomes and function. For example, genomic information can be used to infer the functional capacities [11] and metabolic networks [13, 14] of individual isolates, relate strain-level traits to the dynamics and activities of microbial communities [15], and systematically mine genomes for genes and gene clusters that may encode novel biochemical functions [16–19]. Traditionally, these tasks have been addressed largely independently within their respective fields and often rely heavily on existing genome annotations. As a result, the conventional approaches often have limited generalizability to novel taxa and prediction tasks.

Here, we propose a general framework for characterizing diverse bacterial functional aspects directly from genomic data. Our framework is built on a pretrained foundation model that generates functionally rich embeddings of the genome by contextualizing all protein-coding genes within the full bacterial proteome. This core architecture is then integrated with both unsupervised and supervised models to perform diverse downstream predictive tasks.

Foundation models have achieved notable success in biology. Protein language models such as ESM [20–22] are trained on billions of protein sequences and have been shown to learn embeddings capturing structural and evolutionary features. Genome foundation models often use nucleotide or kmer inputs and focus on relatively short sequence contexts [23–25], making them effective for generating functional representations for small genome segments, but less suited for capturing large-scale genomic features.

Bacterial genomes are more compact and structured than eukaryotes, featuring fewer genes, a much higher proportion of coding sequences, simpler regulatory networks, and local clustering of functionally and metabolically related genes, such as operons and biosynthetic gene clusters [26–30]. Furthermore, although metabolically diverse, many bacterial genes and pathways are conserved across unrelated taxa, often due to horizontal gene transfer [26].

Given these characteristics, we present bacterial proteome transformer (BacPT), an unsupervised model for generating genome embeddings from entire bacterial proteomes (Figure 1A). BacPT was developed using a strategy that efficiently represents a bacterial genome as an ordered sequence of ESM protein embeddings [31– 33] and was trained to generate condensed, context-aware genomic representations through a self-supervised reconstruction objective. This strategy and the smaller gene content in bacteria allows BacPT to take as input all genes in a bacterial genome, in order to model both local and global gene interactions.

**Figure 1.**
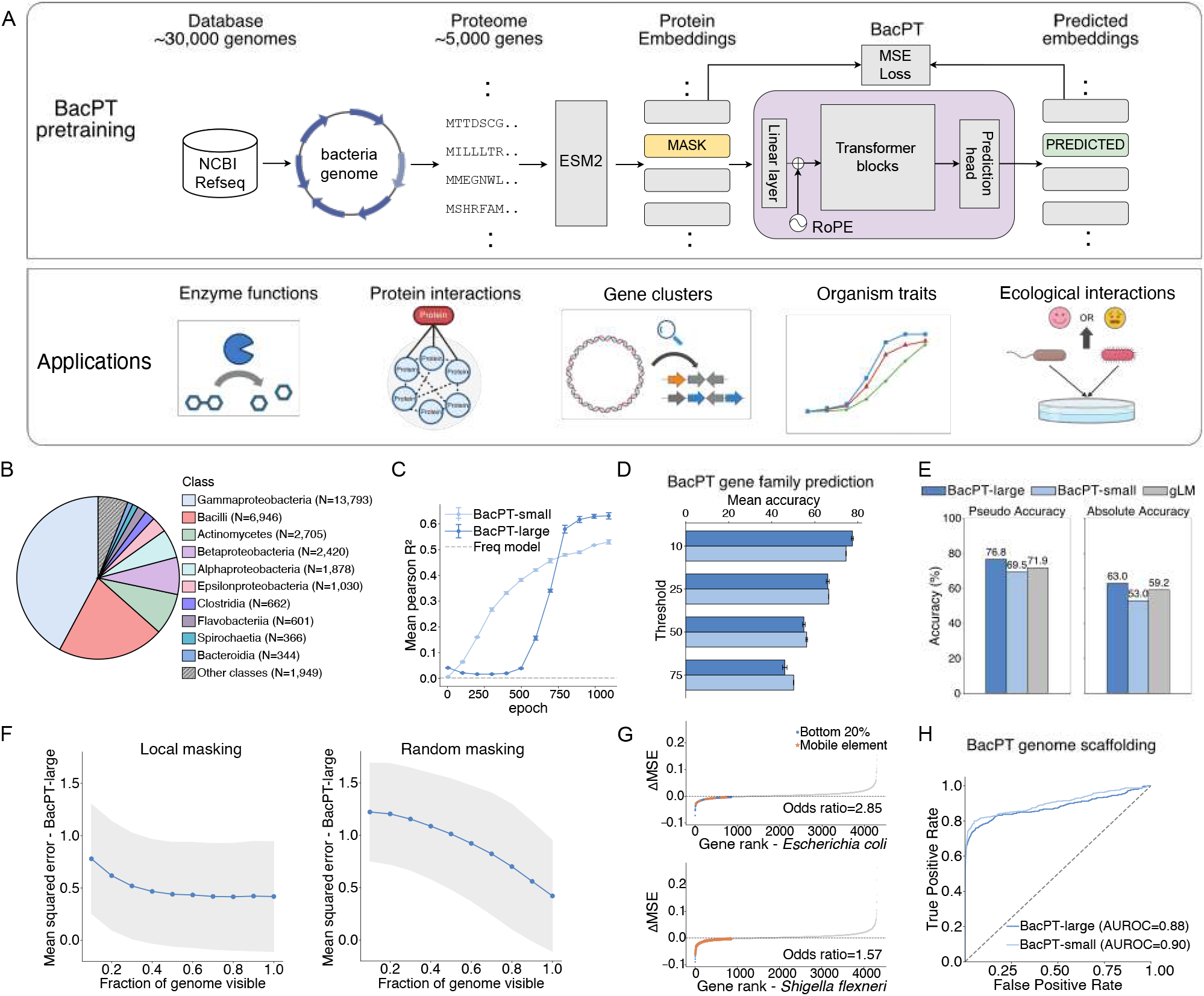
BacPT learns whole-genome contextual information. A. BacPT model architecture and downstream applications. B. Distribution of RefSeq genomes used for BacPT training across different bacterial classes. C. Model performance of BacPT-large and BacPT-small at different training checkpoints. Model performance is measured using squared Pearson *R* between predicted embeddings and input ESM embeddings averaged across randomly masked proteins in 100 test genomes. D. Accuracy of BacPT models at predicting gene family identities for masked proteins. Individual genes were assigned to one of 10,000 gene families, derived by clustering sequences of all proteins in the training data using Faiss-GPU. Test genes were filtered by different percentile rank thresholds to the gene family centroids. E. Absolute and pseudo-accuracy of BacPT models on the test *E. coli* K-12 genome, benchmarked against the metagenomic foundation model gLM. F. BacPT uses whole genome information for gene contextualization, but preferably uses local information. Panels show mean squared error (MSE) of model prediction for single masked proteins when increasing proportion of the bacterial proteome is made visible to the model. Left panel: visible portions of genomes were centered around the masked gene. Right panel: visible portions of the genome were chosen randomly. Results were averaged across 5,059 test genomes. G. Mobile elements are less dependent on local genomic context. ΔMSE values were calculated for individual masked genes when a local genomic window containing 10 genes is made visible to BacPT, compared to predictions where the window is replaced with a random segment from the same genome. Results were calculated using BacPT-large. H. BacPT models can identify correct genome scaffolds. Mean MSE values were calculated for correct whole-genome scaffolds and incorrect genome scaffolds on randomly masked genes. BacPT’s ability to identify correct scaffolds were measured using AUROC, calculated by comparing the mean reconstruction MSE against scaffold labels. Results were aggregated across 100 test genomes.

We trained BacPT on tens of thousands of proteomes derived from high quality RefSeq genomes across diverse bacterial taxa (Figure 1B). We hypothesize that by modeling the probability distribution of natural bacterial proteomes across diverse taxa, BacPT will be able to learn gene covariation due to metabolic and functional coupling, conserved local genomic architecture, and broader gene network organization. This could in turn enable the construction of rich, biologically meaningful latent representations that provide more informative features than raw genomic sequences, uncontextualized protein embeddings [34], or human-curated annotations, making them suitable for a wide range of downstream applications.

We performed analyses of BacPT’s behaviors and showed that it has learned both local and global genomic context. This is further corroborated by benchmarking BacPT’s performance on masked language modeling tasks against previous short context models. Importantly, we demonstrated the utility of the BacPT framework in a number of downstream tasks across different biological scales (Figure 1A). On the single gene level, we found that genomic contextual embeddings enables better prediction of enzyme activity. BacPT can also identify local genomic structures such as operons and biosynthetic gene clusters. We further show that whole proteome representations improves the prediction of organism-level metabolic traits. Finally, we show that the proteome representations can also be leveraged to improve prediction of the outcome of ecological interactions.

## Results

### Unsupervised proteome transformer model for understanding bacterial functional landscapes

To develop a bacterial proteome foundation model, we first curated a dataset consisting of 33,140 complete bacterial genomes downloaded from NCBI RefSeq Genome database [35], representing a diverse array of bacterial taxa (Figure 1B). We chose genomes assembled at the chromosome (or higher) level, to allow our model to learn long-range interactions between genes (see Methods).

We clustered all genomes in our data by applying HDBSCAN [36] on binary vectors representing the presence/absence of 31,096 protein families, which were in turn derived by clustering the amino acid sequences of all proteins in our dataset using MMseqs2 [37] at 50% sequence identity (see Methods). This resulted in three distinct clusters (Supplemental Figure S1). We selected the cluster coincidental with the clade containing the genera *Escherichia, Shigella* and *Salmonella* as our test set (consisting of 5,059 genomes) and used the remaining clusters for training (28,081 genomes; a total of 92.3 million protein sequences). This train-test split strategy ensures the test genomes have minimal similarity to the training set during inference.

For each bacterial genome, we extracted its proteome as the list of all predicted protein coding genes, in their original order in the RefSeq genome. We then use the protein foundation model ESM2 (esm2 t12 35M UR50D8) to generate continuous representations from the amino acid sequences of individual genes in order to encode their functional and evolutionary content (Figure 1A). Since ESM2 produces a 480-dimensional vector for each amino acid within a protein, we averaged these vectors across the entire protein to obtain a single representative vector for each gene.

BacPT adopts a transformer-based architecture that takes as input the mean ESM embeddings of all genes within a genome, projects them through a multi-layer perceptron (MLP), and processes them with a series of transformer layers to model interactions among genes. Through self-attention, the embedding for a gene is updated by integrating information from other genes in the genome. The resulting latent representations in the hidden layers therefore encode both protein features and genomic context, yielding contextualized gene embeddings that can be leveraged for diverse downstream tasks.

We used modified mask language modeling strategies to train BacPT. Specifically, we corrupted input protein embeddings of randomly chosen genes with Gaussian noise and trained BacPT to reconstruct the original ESM embeddings from the genomic context, with the model weights updated using the mean squared error (MSE) loss (Figure 1A). The continuous input and output of our model is in contrast to conventional genome and proteome language models, which predict discrete tokens representing nucelotides [38], k-mers [23], or gene family gitidentities [39]. By operating directly on continuous gene representations, our model preserves the rich biological information distilled by ESM2, which may otherwise be lost through artificial discretization. We developed two models, BacPT-small and BacPT-large, which differ in model complexity, positional encoding schemes, and training strategies. Specifically, BacPT-small utilizes a RoBERTa [40] backbone with a hidden size of 480, 10 transformer layers, and 5 attention heads. It employs relative key-query position embeddings to capture dependencies across sequences of up to 5,000 proteins. In contrast, BacPT-large is built on a RoFormer [41] backbone comprising 19 transformer layers and 10 attention heads. It utilizes Rotary Positional Embeddings (RoPE) to facilitate more robust long-range dependency modeling across the whole-proteome level.

We trained BacPT-large in two stages to capture both local and global genomic features efficiently. In stage one, the model was pre-trained on short genomic contigs containing up to 50 proteins to learn local-level patterns. In stage two, these weights were transferred to the whole-genome scale. This transition enables BacPT to operate at the whole-proteome level for the majority of bacterial genomes. In contrast, BacPT-small was trained on the whole-genome scale across all epochs.

For BacPT-small, we maintained a constant masking rate of 40%. For BacPT-large, we implemented a cosine annealing schedule by reducing the masking probability from 40% at the start of training down to 1% by the final epoch, in order to fine-tune the model to learn richer contextual information.

### BacPT learns whole-genome contextual information

Overall, we observed that the model’s predictive performance on masked genes improved steadily over training epochs, reaching a final genome-averaged *R*^2^ = 0.6 for BacPT-large, and *R*^2^ = 0.5 for BacPT-small, calculated between the input ESM 480-dimensional embeddings and the BacPT-predicted embeddings on randomly masked genes, across 100 random test genomes at 1% masking percentage (Figure 1C; see Methods).

In addition to *R*^2^ between the continuous original and predicted ESM embeddings, we also assessed BacPT’s prediction performance by directly measuring how well BacPT can predict the identity of masked proteins. To this end, we first clustered all proteins in our training data based on their ESM2 embeddings into 10,000 protein families using Faiss-GPU [42] (see Methods). Embeddings of masked proteins are then mapped to the nearest cluster to generate discrete gene family labels. We then calculated prediction accuracy as the proportion of masked proteins whose input ESM2 embeddings and predicted BacPT embeddings are mapped to the same cluster. Overall, we observe that BacPT can adequately predict the family of the masked proteins, with prediction accuracy ranging from 50% to 75% on test genomes, when filtering proteins at varying stringency levels based on their Euclidean distances to the gene family centroids (Figure 1D).

We next benchmarked against gLM [32], a foundation model trained on protein ESM embeddings for bacterial metagenomic contigs. We compared performance of the two models at predicting single randomly masked proteins for the test *E. Coli* K-12 proteome. Performance was measured using two metrics originally designed to gauge the performance of gLM, namely the pseudo accuracy score (frequency the predicted embedding for the masked protein having the smallest Euclidean distance to its original ESM embedding, against all proteins within a local contig of 30 consecutive genes), and the absolute accuracy score (frequency with which the predicted embedding has the smallest Euclidean distance to its original ESM embedding against all proteins within the same genome; see Methods). We found that BacPT-large showed noticeable improvement over gLM (pseudo accuracy = 0.77 vs. 0.72; Figure 1E). The advantage of BacPT-large over gLM also carries over when gauging model predictions using the absolute accuracy score (absolute accuracy = 0.63 vs. 0.59; Figure 1E).

To assess the contribution of local and global genomic context to BacPT, we conducted a detailed experiment where we recorded the prediction performance of BacPT-large on single masked genes in test genomes when different proportions of the genome was made visible to our model (see Methods). We found that the prediction MSE decreases monotonically when increasing fraction of the genome was unmasked (Figure 1F). This indicates that co-variation between genes extends across the entire proteome and that BacPT has learned long-range dependencies through training on complete genomes. Furthermore, we observed that the prediction performance of BacPT-large was higher when the visible genomic portion was centered on the masked protein than when it was randomly sampled across the genome (Figure 1F). This suggests that, although gene interactions extend to the whole genome scale, local gene content is more relevant than long-range context.

Our results so far demonstrate that BacPT is capable of predicting masked proteins from their genomic context. Across a bacterial genome, however, different genes are expected to exhibit varying degrees of dependence on local and global genomic background. For example, mobile genetic elements (such as transposases) can excise and reintegrate themselves into different genomic locations. As a result, compared to core genes, these elements are likely less predictable from their surrounding context alone.

To test this hypothesis, we first identified mobile genetic elements in the *E. Coli* K-12 and *Shigella flexneri* genomes (see Methods). We then used BacPT to predict embeddings for the masked mobile-element proteins both with and without access to a surrounding 10-gene window, and quantified the resulting change in prediction error. We found that the improvement in model prediction for the masked mobile elements attributed to the local window is often negative when measured using reconstruction MSE and ranks among the lowest across all genes in both genomes (Figure 1G), suggesting that BacPT captures genome-scale organizational principles that distinguish context-dependent core genes from highly mobile, context-independent elements.

### BacPT identifies correct genome scaffolding from contigs alone

The results above demonstrate that BacPT models capture both local and global genomic context. This capability suggests that BacPT may enhance bioinformatics applications through its comprehensive understanding of genome structure, such as de novo genome assembly.

In this section, we explore this potential by testing whether BacPT can be applied to bacterial genome scaffolding, which is the problem of ordering and orienting contigs to reconstruct the full chromosome [43]. Traditional scaffolding approaches typically rely on paired-end or long-read sequencing data to bridge gaps between contigs. Here, we investigate whether reasonable scaffolding can be achieved using only contig data.

To this end, we first artificially fragmented 100 test genomes into 2, 3, 5, or 7 contigs of approximately equal sizes, with no sequence overlap between them (Supplemental Figure S2; see Methods). We then used an unsupervised procedure to identify the correct scaffold without additional sources of information. As a result, we must leverage gene co-localization patterns learned during training to infer the correct scaffold.

Our unsupervised procedure uses the genome-averaged reconstruction MSE as a quality measurement for a proposed scaffold, which is calculated over masked genes randomly sampled from the genome scaffold (see Methods). As BacPT is sensitive to both local and global genomic context, it is expected to better predict masked genes under the correct genome scaffold, thereby allowing us to detect incorrect scaffolds.

To test this procedure, we applied BacPT to compute reconstruction MSE for correct genome scaffolds, along with incorrect scaffolds generated by randomly permuting the order and orientation of the contigs. We found that the genome-averaged reconstruction MSE provides a robust metric for distinguishing correct from incorrect scaffolds, with BacPT-large outperforming BacPT-small and achieving an AUROC of 0.9 across 100 random test genomes (Figure 1H).

This result indicates that BacPT embeddings encode sufficient local and long-range information to assess genome-scale structural consistency. This ability to identify correct scaffolding from contigs without orthogonal data makes BacPT a potentially valuable component of larger genome assembly pipelines.

### BacPT gene embeddings improves prediction of enzymatic activities

Having established that BacPT models capture contextual information across whole bacterial genomes, we now perform a series of tasks to assess the utility of BacPT for functional predictions on different biological scales.

We first tested if genome-wide contextualization of single genes can improve the prediction of observable gene functions. To this end, we curated a dataset from the BacDive database [44] containing binary activity profiles for 28 enzymes, with activity measurements derived from standardized API assays (see Methods).

We first examined whether the activities of these enzymes could be predicted simply from the presence/absence of the coding genes. Specifically, we used the contrastive learning model CLEAN [45] to predict the EC number for all genes in a genome and recorded the presence or absence of homologs for all 28 enzymes (Figure 2A). Figure 2B shows the proportion of strains exhibiting measurable enzymatic activity among all strains harboring homologs for the enzyme-coding gene. Interestingly, this percentage varies widely across enzymes, ranging from 10% to 80%, suggesting that presence/absence of the coding gene alone is often an unreliable predictor of enzyme activity.

**Figure 2.**
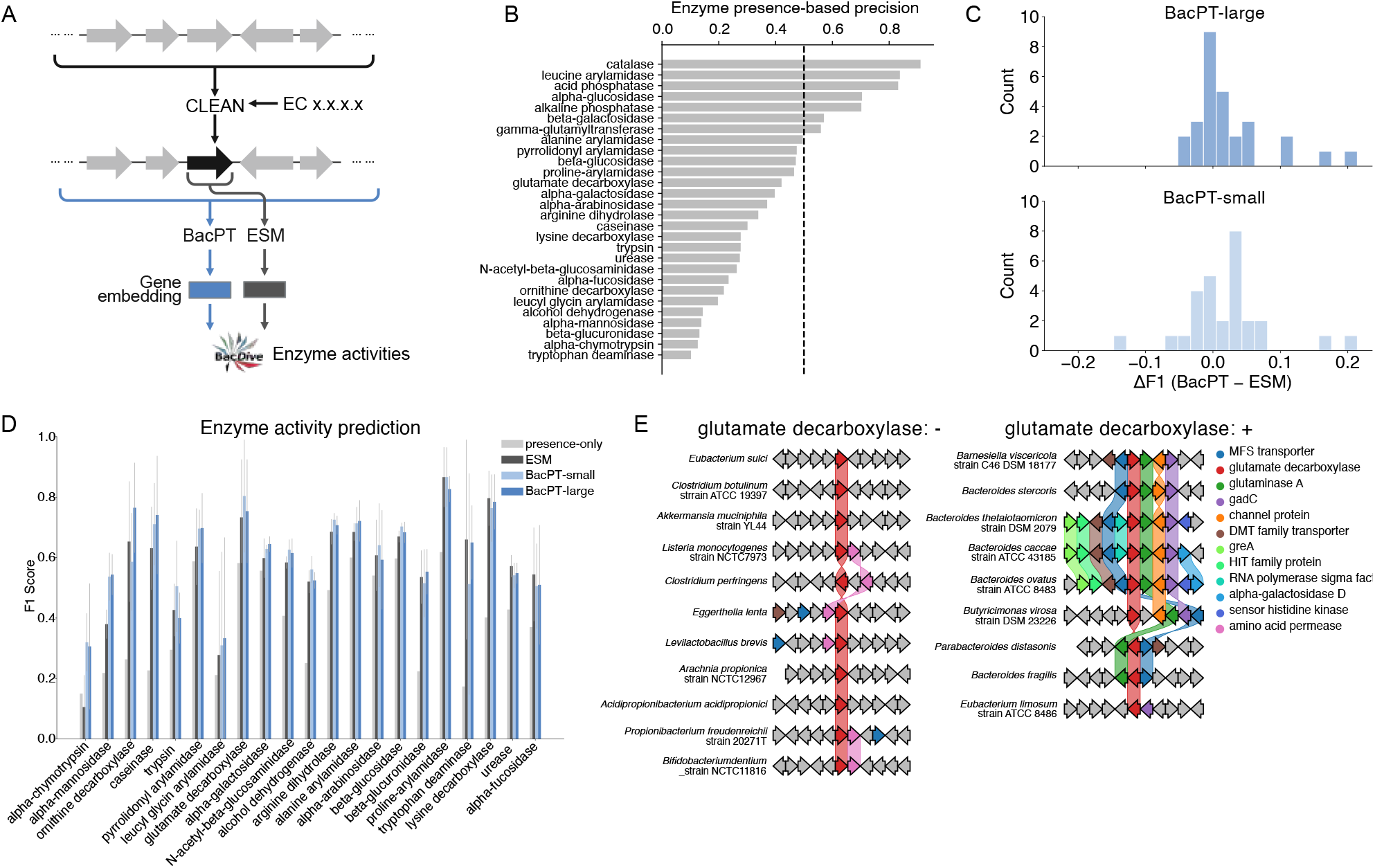
Application of BacPT embeddings for predicting enzyme activities. A. Workflow for predicting activities for 28 enzymes based on the BacDive database. We use the contrastive learning model CLEAN to scan the whole proteome for enzyme-coding homologs using the corresponding EC number as the query. Contextualized embeddings were generated for the identified gene (if present) by passing the whole proteome through BacPT and only retaining the embedding of the focal gene. Uncontextualized embeddings were generated using ESM. Embeddings were then used as inputs to separate linear probes trained to predict binary activity profiles for each enzyme. B. Coding gene presence is not a perfect predictor of enzyme activity. Bars show the proportion of genomes harboring CLEAN-identified homologs that also have enzyme activity. C. Improvement of supervise models trained on BacPT embeddings over models trained on ESM embeddings, measured as Δ*F*_1_ D. Comparison of model performance on enzymes with presence-only precision (Panel C) *<* 0.5. E. Synteny plots for a local 10-gene window centered at the glutamate decarboxylase coding gene for strains with no enzyme activity (left) and strains with activity (right).

Considering that this variation is unlikely the result of annotation errors alone given the previously demonstrated high prediction accuracy of CLEAN [45], it may instead reflect true functional divergence among homologs or strong dependency of the enzyme-coding genes on their genomic backgrounds. To test this hypothesis, we applied a supervised learning approach and trained logistic regression models to refine the prediction of enzyme activity while restricting to strains harboring the coding gene (see Methods). Our baseline model was trained on the raw 480-dimensional ESM embeddings averaged across all amino acid positions of the coding gene (Figure 2A).

To test if we can further improve performance by enriching ESM embeddings with genomic context, we retrained logistic regression models on the 480-dimensional BacPT embeddings of the focal enzyme-coding gene, contextualized using all genes in the genome (Figure 2A). Overall, linear probes trained using BacPT-contextualized embeddings achieved higher prediction accuracy than models trained on uncontextualized, raw ESM gene embeddings (Figure 2C), with BacPT-large showing overall stronger improvement.

Among enzymes with fewer than 50% of strains exhibiting positive activity conditional on gene presence, there are substantial variation in performance gains, with the largest improvements observed for alpha-chymotrypsin, alpha-mannosidase, ornithine decarboxylase, and caseinase (Figure 2D). Additionally, we also found that linear probes based on ESM embeddings showed higher F1 scores than predictions made solely based on gene presence (Figure 2D), suggesting that evolutionary and functional information encoded in the ESM embeddings for the enzyme-coding genes also contribute meaningfully to variations in enzyme activity, in addition to the genomic context.

To understand how genomic context influences enzyme activity, we provide a close examination of the local neighborhood of the gene glutamate decarboxylase. First, genomes lacking activity showed little to no conserved organization within a window of ten genes (Figure 2E). In contrast, we found strong synteny in genomes for strains with measurable enzyme activity. In particular, functional glutamate decarboxylase is frequently co-localized with genes including glsA, MSF transporter, DMT family of potassium transporters, gadC, which can influence the activity of glutamate decarboxylase by modifying cellular aspects such as substrate availability, product removal, pH, ion balance, and transport dynamics.

Overall, our findings suggest that genomic context can indeed play important roles in determining the enzymatic activity of a gene. Furthermore, BacPT can generate meaningful genome-contextualized gene embeddings, which can be exploited for improving gene function predictions.

### BacPT captures gene interactions and gene clusters

Results from the previous section show that BacPT can improve enzyme activity prediction by contextualizing the focal genes using the local genomic region. In this section, we test whether the model can also be used to directly identify different types of gene clusters, which will make it a useful tool for annotating poorly studied genomes and discovering novel gene clusters.

As a first step, we use a supervised approach to test if BacPT embeddings can be used to identify operons, which are conserved DNA segments containing clusters of adjacent genes that are co-regulated and co-transcribed. We curated a dataset using RegulonDB [46], consisting of operons containing more than nine genes in the *E. coli* K-12 genome. We found that a logistic regression model trained on mean ESM embeddings of all genes within the operonic window achieved an accuracy of 0.66 and an AUROC of 0.73 (Figure 3A; see Methods), when evaluated on test data consisting of positive operonic and size-matched non-operonic windows. This suggests that certain protein features are likely overrepresented in operons, thereby allowing the ESM2-based model to make non-trivial predictions.

**Figure 3.**
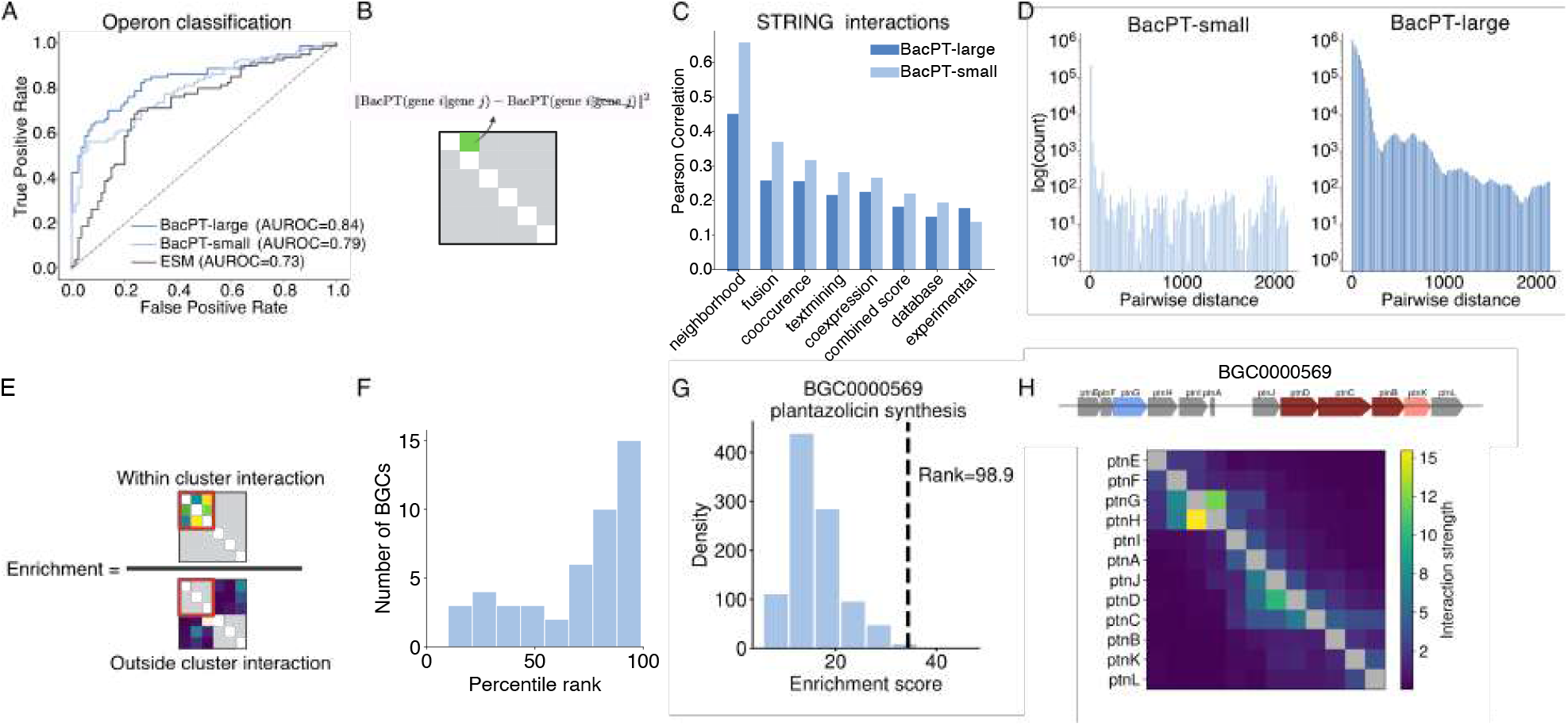
Application of BacPT for identifying gene clusters. A. Performance of supervised models at identifying operons in the *E. coli* K-12 genome. Models were trained using BacPT-small, BacPT-large, or uncontextualized ESM embeddings. B. Jacobian metric for quantifying gene interactions. For gene *i*, its embedding is first replaced with noise. Squared norm of the difference between predicted embeddings for gene *j* with and without gene *i* is used to quantify the influence of gene *i* on gene *j*. C. Correlation of the Jacobian matrix against various interactions in the STRING database for the *E. coli* K-12 genome. D. Distribution of distance (measured as number of intervening genes) for the top 1 % of interacting gene pairs. Distance for gene pair (*i, j* was calculated as the minimum number of genes positioned between the gene *i* and *j*. E. Enrichment score for a gene cluster (red square) is equal to the ratio between the median interaction strength for genes within the cluster and median interaction strength for genes within the cluster and genes outside the cluster. F. Percentile rank distribution for 82 biosynthetic gene clusters (BGCs) from 8 genomes. G. BGC0000569 is enriched for interactions among genes within the cluster. H. Interaction strengths between genes within BGC0000569. Shown is the diagonal block for genes in this BGC with entries given by the Jacobian values.

Importantly, the linear probes trained on BacPT-contexualized embeddings showed noticeable improvement over the performance of the ESM model, with BacPT-large achieving an accuracy of 0.75 and an AUROC of 0.84 (Figure 3a). This suggests that our model has learned some general high-level principles of local gene coupling.

Next, we set to test an unsupervised framework for identifying gene clusters and interaction networks. To this end, we first generated an all-by-all interaction matrix *M*, where the directional interaction score *M*_*i,j*_ for proteins *i* on *j* is calculated as the *L*_2_ norm of the difference in predicted embeddings for protein *j*, when the input ESM embeddings for gene *i* is replaced with a zero vector (see Methods; Figure 3B). *M*_*i,j*_ will approximate zero, if *i* and *j* represent random pairs of proteins, as *i* provides little information for predicting *j*. If instead *i* and *j* tend to co-occur across many genomes in the training data, we expect BacPT to have learned this relationship, resulting in higher *M*_*i,j*_ values. Thus, *M* provides a measurement of statistical interaction strengths between genes within a genome, similar to how the categorical Jacobian [47] quantifies pairwise interaction between amino acids within proteins.

We first observed that the distribution of interaction scores is highly skewed in the *E. Coli* K-12 genome, with most gene pairs having low scores (Supplemental Figure S3). We also found that the majority of strongly interacting genes are typically located in close adjacency, although strong interactions over long physical distances are also present (Figure 3D).

To test the types of gene interactions enriched in our Jacobian matrix, We compared *M* with various gene interaction statistics derived from the STRING database [48]. We found that the symmetrized version of *M* (see Methods) correlates the strongest with neighborhood interactions in STRING, with BacPT-small showing stronger correlation (Pearson *r* = 0.65) than BacPT large (Pearson *r* = 0.45; Figure 3C). We also observed moderate correlation between *M* and STRING interactions including fusion, co-occurrence, and co-expression scores. Furthermore, *M* also shows small but significant correlation (*r >* 0.15 for both models) with experimentally measured protein interaction strengths (Figure 3C).

This findings confirm our earlier result that BacPT preferentially leverages local-level information when generating genome-contextulized embeddings. Therefore, this ability to identify co-localizing genes potentially make BacPT a tool for identifying gene clusters in unannotated genomes [18, 19]. To assess the feasibility of this idea, we applied the BacPT Jacobian matrix to 82 biosynthetic gene clusters from 8 species with ≥5 genes, derived from the Minimum Information about a Biosynthetic Gene Cluster (MiBIG) database [49] (see Methods). Specifically, for each gene cluster, we calculated an enrichment ratio using the Jacobian matrix derived from BacPT-small equal to the median interaction strength between genes within the cluster and median interactions between genes in and outside the cluster (Figure 3E). Overall, we find that enrichment scores for BGCs exhibit high percentile ranks in null distributions of size-matched non-BGC clusters (Figure 3F), suggesting that that BGCs tend to harbor more statistically coupled genes. For example, the 12 genes in the BGC for plantazolicin synthesis (MiBIG id: BGC0000569) show high enrichment against the genomic background (percentile rank=98.9; Figure 3G), evidenced by a number of strong and moderate interactions (Figure 3H).

Overall, our exploratory analyses confirm that BacPT has learned local gene co-variation. The utility of zero-shot procedures and simple classifiers trained on BacPT embeddings indicates that BacPT may serve as a strong foundation model for studying gene interactions and identifying gene clusters in bacterial genomes.

### BacPT improves bacterial metabolic trait prediction

Having established that BacPT generates biologically rich embeddings at the gene level and effectively decodes the local syntax of the bacterial genome, we next examine whether the contextualized gene representations can be aggregated to capture higher-level functional aspects of the entire bacterial cell.

Specifically, we trained a series of supervised models using BacPT embeddings to predict different metabolic phenotypes (Figure 4A). The input to the supervise model consists of mean BacPT embeddings across all genes, which is then fed through a simple linear probe to predict phenotypes (see Methods).

**Figure 4.**
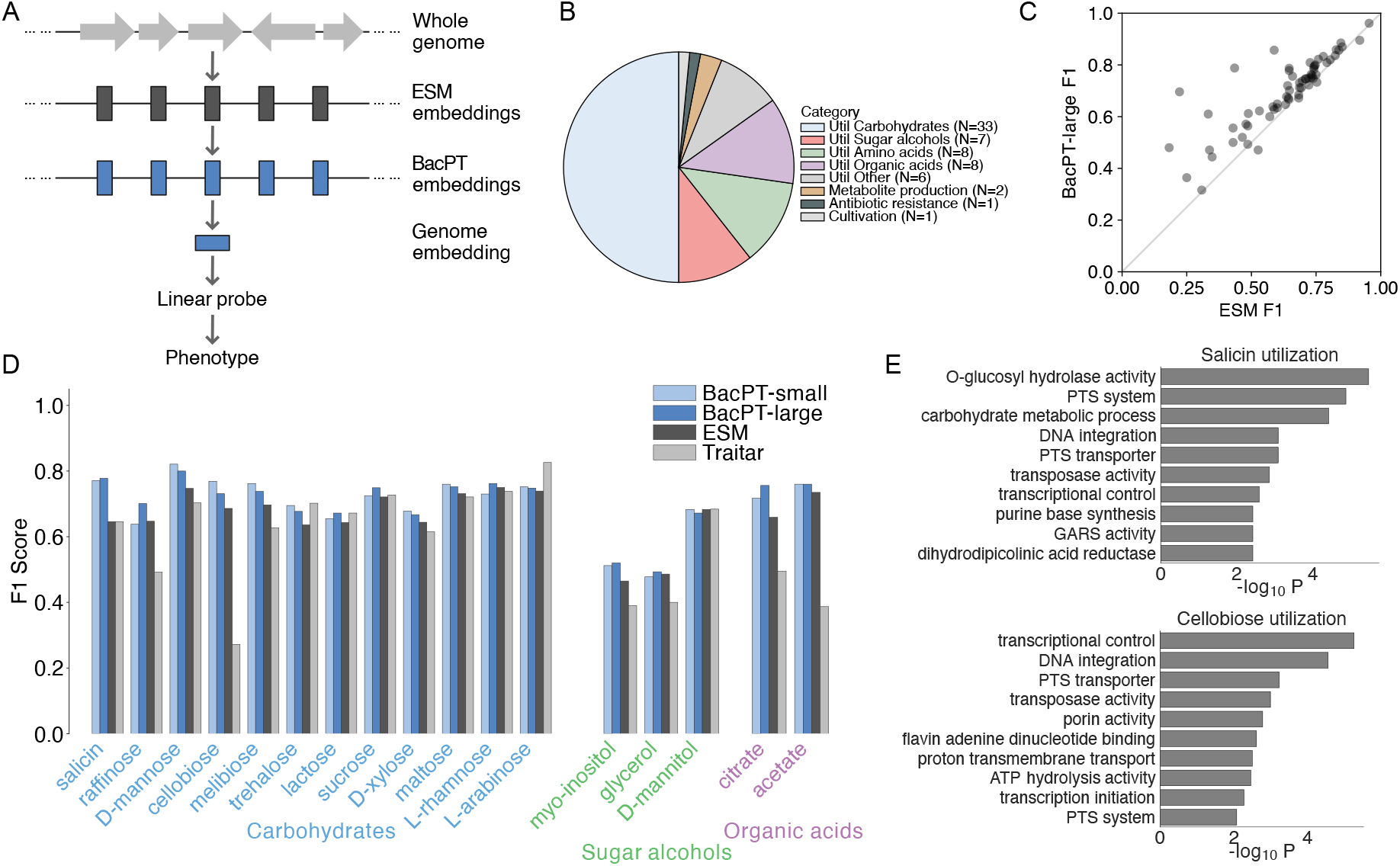
BacPT enables better bacterial trait prediction. A. Architecture of supervised models for predicting bacterial phenotypes from BacPT-contextualized embeddings. B. Distribution of BacDive-derived traits. C. Performance comparison between linear probes trained on BacPT embeddings and ESM embeddings. D. Detailed performance comparison of linear probes trained using BacPT and ESM embeddings against Traitar on 17 shared metabolic traits. E. Gene Ontology (GO) term enrichment in the top 500 Pfam identifiers discovered by applying perturbation-based attribution analysis to BacPT-based linear probes for salicin and cellobiose utilization.

Phenotypic data were retrieved from the BacDive database and mapped to RefSeq genomes using shared accession numbers (see Methods). We included all available cellular phenotypes but excluded enzymatic activities, as these were analyzed in the preceding section. To ensure statistical robustness, we retained only traits with sufficient class balance and a minimum sample size of 10 per class (see Methods). This resulted in a final dataset of 66 traits, including 62 phenotypes profiling the utilization of various metabolites (Figure 4B), with additional phenotypes covering biosynthesis, antibiotic resistance, and growth conditions.

To test if the genome-contextulized embeddings generated by BacPT better predict these organismal phenotypes, we first compare the performance of linear probes trained on genome-averaged BacPT embeddings against models trained using mean ESM gene embeddings (see Methods). As shown in the scatter plots (Figure 4C), BacPT-large consistently outperforms the uncontextualized ESM baselines across nearly all tested traits (Figure 4C), with median improvement in F1 = 7%. Similar trend also holds for linear probes trained using BacPT-small embeddings (Supplemental Figure S4).

To further assess the utility our foundation model-based framework for trait predictioin, we also benchmarked linear probes based on BacPT and ESM embeddings against Traitar [50], a microbial phenotype prediction method that works by mapping genomic protein families (Pfam) to phenotypic traits through a pre-trained manual annotation pipeline. Across 17 traits predicted by both BacPT and Traitar, we found that supervised models trained on ESM and BacPT embeddings outperform Traitar across different categories of metabolites, with BacPT models further exhibiting moderate but consistent improvement over ESM models (Figure 4D).

Specifically, the contextualized representations from BacPT yielded more accurate predictions, most notably for cellobiose, where BacPT achieved a maximum of 49.7% improvement over Traitar and 8.3% over ESM, citrate (26.1% over Traitar and 9.7% over ESM), and salicin (13.2% over Traitar and 13.2% over ESM). In addition to benchmarking against supervised ESM model, we also compare our model performance against a structured supervised model that uses presence/absence of Pfam domains in a genome to predict phenotypes. Briefly, presence/absence data is generated using InterProScan [51], resulting in a 8,552-dimensional binary vector. We find that BacPT-based supervised models maintain a moderate performance margin over the structure Pfam model (mean 3% improvement in F1 by BacPT-large; Supplemental Figure S5).

To further demonstrate that BacPT-based models can be interpreted to distill biological insights, we next performed a perturbation-based attribution analysis using the supervised models trained with BacPT embeddings to understand the genetic determinants contributing to phenotype predictions. Because the input to BacPT consist of unstructured proteomes represented as ESM embeddings, we first annotated all test genomes with Pfam domain identifiers using InterProScan, to allow gene-level attribution scores to be aggregated across test genomes at the domain level (see Methods). For each genome, we systematically perturbed the embeddings of genes associated with a given Pfam domain by replacing them with zero vectors to effectively remove their information content and measured the resulting change in prediction. Domain attribution was quantified using the metric Δ_total_, equal to the difference between baseline and perturbed logits (see Methods).

To assess the biological relevance of the identified domains, we performed Gene Ontology (GO) enrichment analysis on the 500 top-ranked Pfam identifiers based on their mean attribution scores Δ_total_. Overall, we find that Pfams with strong effects on trait prediction generally are enriched for trait-relevant GO terms.

For salicin, the enrichment analysis identifies terms directly associated with the degradation of complex glycosides (Figure 4E). Notably, O-glycosyl hydrolase activity corresponds to the enzymatic cleavage required for salicin utilization. Phosphoenolpyruvate-dependent sugar phosphotransferase (PTS) system is a carbohydrate transport system found in most bacteria for translocation of sugar, in particular salicin [52]. GO analysis also reveal other transport and catabolic machinery, such as general carbohydrate metabolic processes and transcriptional regulation.

For cellobiose, the result highlights a focus on genomic regulation and transport (Figure 4E). The topranked term is the regulation of DNA-templated transcription, followed by DNA integration. These findings suggest that the domains most predictive of cellobiose utilization are often found within regulated operons or mobile genetic elements. Similar to salicin, PTS system also appears to have strong effects, highlighting the importance of PTS-mediated uptake for this disaccharide.

Overall, we have shown that BacPT-contextulized gene embeddings can be aggregated to represent diverse functional aspects of the whole bacterial genome, which can be exploited to predict organismal phenotypes. Furthermore, we showed that attribution analysis can be applied to specialized supervised learning models for understanding the genetic basis of metabolic traits.

### BacPT improves prediction of ecological interactions

The previous section establishes that BacPT embeddings are effective for predicting organismal phenotypes. Because ecological interactions between bacteria in natural environments are largely trait-mediated [53, 54], we hypothesize that the genomic functional aspects captured by the contextualized BacPT embeddings useful for predicting metabolic phenotypes may also be extended beyond the organism level to predict ecological interactions.

To test this, we applied BacPT embeddings to an ecological interaction dataset consisting of pairwise interaction outcomes among 20 species across 40 nutrient conditions [55] (see Methods). For each environment, we trained a linear probe using the concatenated mean BacPT embeddings of the two species to predict interaction outcomes (mutualism, competition, or parasitism) (Figure 5A; see Methods). For benchmarking, we trained analogous models using the genome-averaged ESM embeddings of the interacting species.

**Figure 5.**
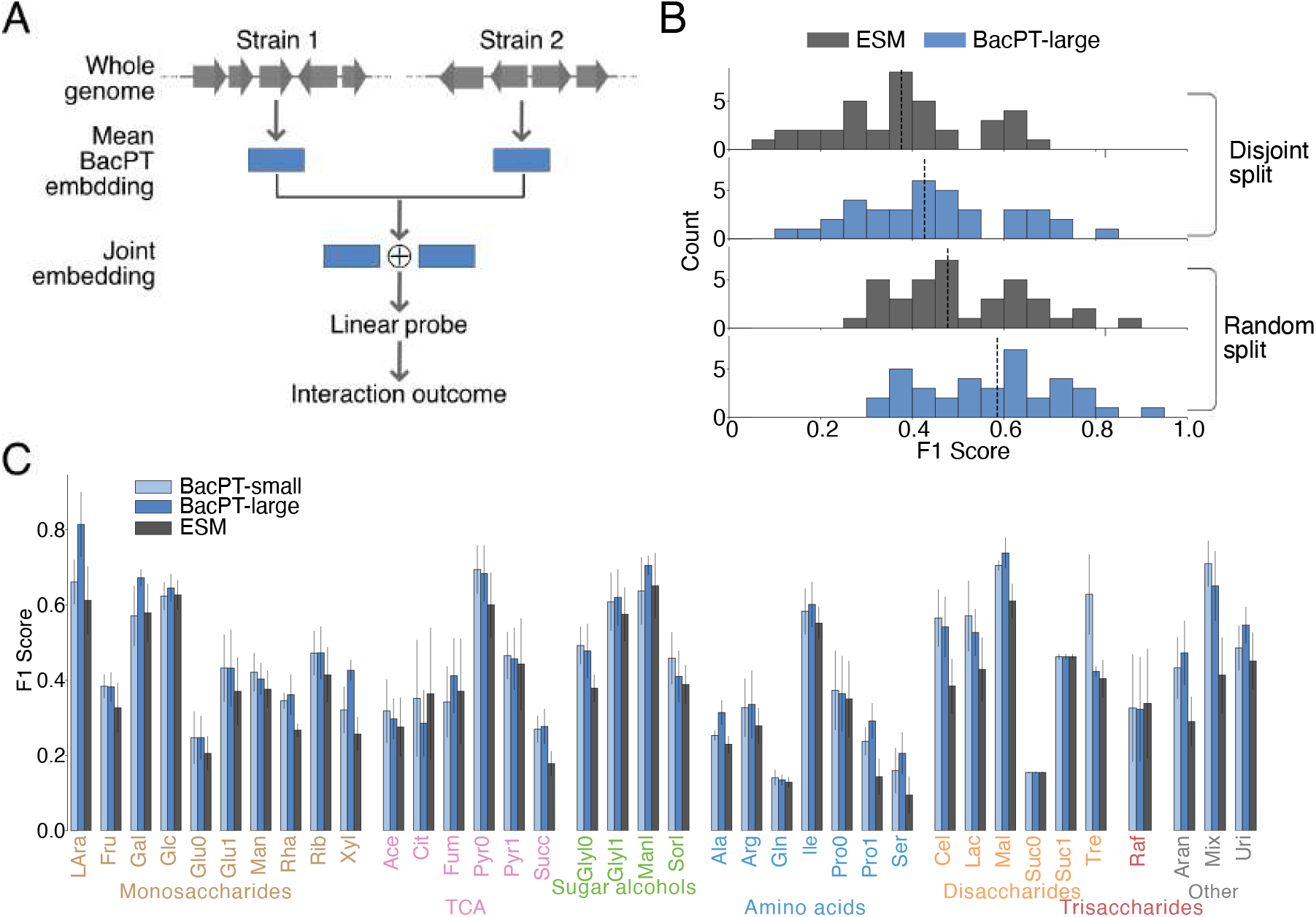
Application of BacPT embeddings for modeling ecological interactions between bacterial strains. A. Workflow for training supervised models for predicting ecological interaction outcomes. Mean genomic embeddings of the two interacting strains were generated by applying BacPT to the whole proteomes. The mean embeddings are concatenated and fed through a linear probe to predict relationship (mutualism, competition, or parasitism) under certain nutrient conditions. B. Distribution of macro-averaged F1 scores of linear probes trained on BacPT embeddings and ESM mean embeddings in 37 carbon sources, under disjoint and random train-test split. Note that prediction performance improves for random split, where test interactions were randomly sampled from all pairs of species. C. Direct comparison of performance of linear probes trained using ESM embeddings and BacPT embeddings, in different carbon environments. Model performance was evaluated under the disjoint train-test split strategy. Suffix 0 and 1 correspond to concentration of the carbon source at 0.5 percent and 0.05 percent, respectively.

To assess model robustness and generalizability, we employed two distinct train-test split strategies. In ‘disjoint split’, the total pool of species was partitioned into mutually exclusive sets; any species present in the test set was completely absent from the training set. In ‘random split’, interactions in the training set were sampled randomly from the total pool of possible pairs. Thus, disjoint split represents a more stringent criterion as the model must distill more general functional ecology principles in order to infer interaction outcomes for pairs involving only unseen species.

First, for disjoint split, we found that supervised models trained using BacPT embeddings overall outperform models trained using mean ESM embeddings, with a mean 7% improvement in macro-averaged F1 score (Figure 5B). Next, when assessing model performance using the random split strategy, we observed an overall improvement in model performance compared with disjoint split, with BacPT models maintaining a similar performance margin (Figure 5B).

In Figure 5C, we show the detailed model performance across 37 carbon sources under the disjoint split strategy. First, we observed great variability in model performance across carbon sources. This could indicate that the genetic architectures of different metabolic pathways vary in their complexity, where some substrates possess more conserved genomic signatures that models can easily identify, while others involve more diverse enzymatic repertoires and stronger dependence on the genetic background.

Notably, we see that BacPT models outperform baseline ESM models at predicting interaction outcomes in all carbon source categories. For monosaccharides, the largest gains by BacPT model were seen for arabinose, rhamnose, and xylose, where models based on BacPT-large embeddings achieve 10-20% improvement in marco-averaged F1 over ESM models. Improvement is also evident in nutrients supporting central carbon metabolism, with gains observed for both TCA-associated substrates and sugar alcohols, including pyruvate (8.3%) and glycerol (9.93%). Among amino acid conditions, the most notable gain was seen for serine (11.09%). In disaccharides, substantial improvements were observed for cellobiose (15.73%) and maltose (12.76%). Across all environments, the highest improvement was observed under mixed carbon conditions, where BacPT-large model outperforms the ESM model by 23.73%.

Taken together, these results demonstrate that the functionally meaningful genomic signals captured by BacPT embeddings are also informative on the level of ecological interactions. The consistent improvement over ESM-based models across diverse nutrient conditions and different train-test split strategies suggest that BacPT models have learned generalizable principles of interaction rather than relying on species-specific rules. This highlights the potential of a unified framework for building predictive models bridging genomic information of microbial communities and ecological functions.

## Discussion

We have developed BacPT, a foundation model trained on 33,140 full-size bacterial proteomes. By representing genomes as ordered sequences of protein ESM embeddings, BacPT captures the complex genomic architecture of diverse bacterial taxa. We demonstrated that BacPT embeddings can be successfully applied across biological scales, ranging from single-enzyme activity to organism-level metabolic traits, and even pairwise ecological interactions. This framework provides a powerful, general-purpose approach for characterizing bacterial functional traits directly from genomic data. And compared to traditional pipelines, BacPT represents a generalizable strategy that can be applied to novel taxa and tasks without heavy reliance on existing human-curated annotations.

Our analyses reveal that BacPT is able to use the whole genome to contextualize individual genes and has learned higher-order structures such as gene interactions and gene clusters. This ability to to encode diverse functional aspects of the bacterial genome is evidenced by the overall superior performance of BacPT-based supervised models over models trained using ESM embeddings. A key distinction arises when comparing supervised models trained on BacPT embeddings to models based on raw ESM embeddings: while ESM-based models primarily capture the additive effects of individual proteins, BacPT-derived representations account for the statistical interactions between genes through genome contextulization. Consequently, these two approaches can be understood as representing additive and epistatic models [56–64] of bacterial function, respectively. Importantly, these epistatic interactions are not learned for specific downstream tasks; rather, they are captured during the self-supervised pretraining phase. This demonstrates that the contextualized embeddings are sufficiently general to serve as a robust foundation for a wide array of prediction tasks. In future applications, the framework could be further improved by learning task-specific gene contextualization, to potentially enable finer resolution in modeling complex biological phenotypes.

While using protein embeddings of all coding genes provides a concise and comprehensive representation of the genome, the integration of regulatory information in non-coding regions represents an opportunity to further refine our model. Although the vast context length of full bacterial genomes may pose a significant technical challenge for traditional transformer architectures, recent breakthroughs in long-context modeling (such as the Evo 2 architecture [65]) may be helpful in bridging this gap.

Our current framework for predicting bacterial functional traits largely relies on labeled data to train task-specific supervised models. This requirement restricts the applicability of BacPT to well-characterized phenotypes and limit our ability to deploy this framework for poorly studied or novel traits where experimental labels are scarce. To overcome these barriers, future developments may investigate more sophisticated pretraining strategies to better tailor the BacPT base models for diverse downstream applications. Additionally, integration of zero-shot or few-shot prediction strategies [66, 67] may allow us to generate accurate inferences in data-sparse regimes and reduce the dependency on comprehensive experimental datasets, thus may be instrumental in broadening the scope of our current framework.

Beyond prediction tasks demonstrated in this paper, our framework may be expanded to encompass other applications. For example, our results suggest that BacPT is well-suited for diverse bioinformatics tasks including genome scaffolding and automated gene function annotation, and may thus be integrated into existing pipelines to boost performance. Furthermore, the ecological interaction tasks may be generalized to predict the functions of entire bacterial communities, such as synthetic communities and microbiomes [68, 69]. As more experimental data and annotations become available, we expect the range of applications for BacPT will continue to expand.

## Methods

### Curation of BacPT training data

Our dataset comprised protein sequences from 33,140 bacterial species downloaded from the NCBI Genome database [35], representing diverse taxonomic families. To ensure minimal similarity between training and test genomes, we performed clustering analysis at both the protein and genome levels. Proteins across all genomes were first clustered using MMseqs2 [37] at 50% sequence identity to define 33,204 protein families. Each genome was then represented as a binary vector indicating the presence or absence of each protein family. Genomes were subsequently clustered using HDBSCAN [36] based on these binary representations to group genomically similar species. Since HDBSCAN is a noise-aware clustering algorithm, genomes classified as noise were removed from the dataset. We designated the cluster containing the genera *Escherichia, Shigella*, and *Salmonella* as the test set (5,059 genomes) and used all remaining clusters for training (92.3 million protein sequences with possible redundancy).

Protein embeddings for all predicted genes were generated using ESM2 (esm2 t12 35M UR50D8), which produces a 480-dimensional vector for each amino acid within a protein. These vectors were averaged across the full protein sequence to obtain a single representative 480-dimensional embedding per gene, such that each bacterial genome is represented as a sequence of 480-dimensional vectors, one per protein.

### BacPT model architecture and training

We trained two models, BacPT-small and BacPT-large, which differ in their architectural backbone and training strategy. BacPT-small uses a RoBERTa [40] backbone with relative key-query position embeddings, while BacPT-large uses a RoFormer [41] backbone with Rotary Position Embeddings (RoPE). BacPT-large was trained in two stages to enable efficient learning across full-length bacterial genomes.

We adapt the RoBERTa architecture for our smaller model, BacPT-small, which has a hidden size of 480, 10 transformer layers with 5 attention heads, and relative key-query position embeddings to capture positional dependencies across sequences of up to 5,000 proteins. BacPT-small was pre-trained using a strategy similar to masked language modeling. Since our inputs comprise continuous embeddings rather than discrete tokens, we progressively corrupted the targeted embeddings with random noise instead of replacing them with a constant vector. Specifically, we applied noise to 40% of proteins randomly selected from a genome and trained the model to reconstruct the original embeddings using mean squared error (MSE) as the loss function. We kept the fraction of noised proteins at 40%, since it has been shown to outperform the masking fraction of 15% for large-size BERT models [70]. As training progressed, the noise magnitude was increased until the masked proteins were completely replaced by Gaussian noise. BacPT small was pre-trained for 1,100 epochs on 16 NVIDIA A100 GPUs.

BacPT-large was trained in two stages using a RoFormer backbone with RoPE, comprising 19 hidden layers, 10 attention heads, a hidden dimension of 480, GELU activations, and a dropout probability of 0.1. In the first stage, a contig model was pre-trained on short genomic contigs of maximum length 50 proteins. At each training step, 15% of protein embeddings were randomly selected and replaced with Gaussian noise sampled with zero mean and variance matching the empirical variance of the training embeddings: ***ϵ***∼ 𝒩 (**0**, *σ*^2^**I**), where *σ*^2^ is estimated from the training data. The model was trained to reconstruct the original embeddings at masked positions using MSE loss. Training was conducted for 500 epochs with a batch size of 1,024, a constant learning rate of 1 × 10^−5^ with 500 linear warmup steps, and an AdamW optimizer with weight decay of 0.1, using 99.95% of the data for training and the remaining 0.05% for validation.

In the second stage, the contig model weights were transferred to initialize a full-length model supporting genomes of up to 5,000 proteins. Since the RoPE position embeddings were trained on sequences of length 50, they were extended to length 5,000 via linear interpolation before being loaded into the new model, while all remaining weights were transferred directly. The full-length model was then pre-trained for 1,100 epochs with a batch size of 8 and a constant learning rate of 1 × 10^−5^ with 500 warmup steps, using 98% of data for training and remaining data for validation. To progressively increase task difficulty over training, the masking probability was annealed via a cosine schedule from 40% at the start of training down to 1% by the final epoch:

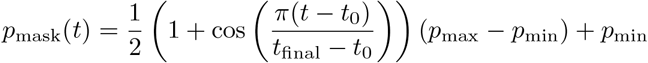

where *p*_max_ = 0.4, *p*_min_ = 0.01, *t*_0_ is the starting epoch, and *t*_final_ is the final epoch, similar to [71]. At each epoch, selected protein embeddings were replaced with Gaussian noise sampled from ***ϵ*** ∼ 𝒩 (**0**, *σ*^2^**I**), and the model was trained to reconstruct the original embeddings using MSE loss. Both stages were trained on 16 NVIDIA B200 GPUs.

### Model accuracy

To assess reconstruction quality over the course of training, BacPT was evaluated at regular checkpoint intervals (every 99 epochs from epoch 3 to epoch 1100) on a random subset of 100 test genomes per replicate, repeated across three replicates. At each checkpoint, 1% of protein embeddings were randomly selected and replaced with Gaussian noise (masked), and the model was used to predict the original embeddings at masked positions. Reconstruction quality was measured using the mean Pearson *R*^2^ between the predicted and ground truth embeddings, computed per genome as:

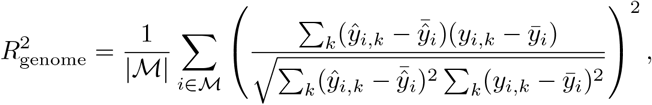

where ℳ is the set of masked positions, *ŷ*_*i,k*_ and *y*_*i,k*_ are the predicted and ground truth embedding values at position *i* and dimension *k* respectively. Mean *R*^2^ and its standard deviation were reported across three replicates at each checkpoint. As a baseline, a frequency model was constructed by replacing each masked embedding with a randomly sampled embedding drawn from a random protein in a random training genome, and the same Pearson *R*^2^ metric was computed against the ground truth embeddings.

Reconstruction accuracy was additionally evaluated on the full *E. coli* K-12 genome (4,297 proteins). Each protein position was masked iteratively by replacing its embedding with Gaussian noise while keeping all other protein embeddings intact, and BacPT was used to predict the masked embedding using the full genomic context. Two accuracy metrics were computed from the resulting set of predicted embeddings to facilitate comparison between BacPT models and the metagnomic foundation model gLM [32]. Absolute accuracy assesses whether the predicted embedding for a masked protein was closest, in Euclidean distance, to the ESM embedding of that same protein among all 4,297 proteins in the genome:

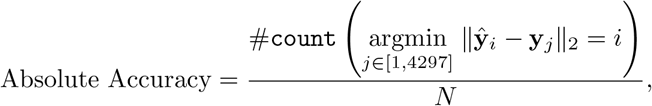

where *N* = 4,297 is the total number of genes in *E. coli* K-12. Pseudo accuracy relaxes this criterion by restricting the candidate set to a local subcontig of 30 consecutive proteins, assessing whether the predicted embedding was closest to the correct protein within that local neighborhood:

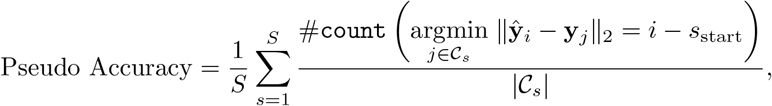

where *S* is the total number of non-overlapping subcontigs of length 30, 𝒞_*s*_ denotes the set of protein indices in subcontig *s*, and *s*_start_ is the starting index of subcontig *s*. Pseudo accuracy thus provides a more lenient measure of reconstruction fidelity by evaluating whether the model correctly identifies a masked protein within its local genomic neighborhood.

### Gene family prediction

Both BacPT-large and BacPT-small were additionally evaluated on the test set using a cluster assignment accuracy metric. All proteins in the training data were clustered into 10,000 clusters based on their ESM2 embeddings using Faiss-GPU [42], with 20 iterations of k-means. For evaluation, 15% of protein embeddings in each test genome were randomly masked by replacing them with Gaussian noise, and BacPT was used to predict the embeddings at masked positions. Both the predicted embeddings and the original ESM2 embeddings were independently assigned to their nearest cluster using the trained k-means index. A predicted embedding was considered correctly assigned if it was mapped to the same cluster as its corresponding ESM2 embedding. To focus evaluation on well-represented clusters, only proteins whose nearest-cluster distance fell below a given percentile threshold were included in the accuracy computation. Accuracy was computed across distance thresholds at the 10th, 25th, 50th, and 75th percentiles, and reported as the mean over three replicates, each evaluated on a random subset of 100 test genomes.

### Effects of masking proportion on prediction accuracy

To assess the how local and global genomic context contribute to BacPT predictions (i.e. results shown in Figure 1F), we performed exploratory analysis of the behavior of BacPT-large. Specifically, we examine how increasing the fraction of genomes visible to BacPT-large affects the reconstruction accuracy for a focal protein. The protein at the genomic midpoint was designated as the focal gene. Its embedding was replaced with Gaussian noise, while the model was provided embeddings for an increasing fraction *f* ∈ 0.1, 0.2, …, 1.0 of the remaining genome as context, with all positions outside the context window zeroed out. Two context selection strategies were evaluated across all genomes in the held-out test set. In the local strategy, context proteins were drawn from a contiguous window centered on the focal gene, spanning positions [ ⌊*N/*2⌋ − *f N/*2, ⌊*N/*2⌋ + *f N/*2], where *N* is the genome length. In the random strategy, the same fraction *f* of proteins were selected uniformly at random from across the genome, without any spatial constraint relative to the focal gene. This design allows direct comparison between the utility of local genomic neighborhood versus dispersed genomic context for reconstructing a masked gene. BacPT was then used to predict the masked focal gene embedding under both strategies, and reconstruction quality was measured as the MSE between the predicted and ground truth embeddings: 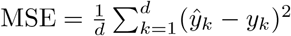, where *d* = 480 is the embedding dimension. Mean MSE and its standard deviation were reported across all test genomes for each context fraction.

### Mobile elements

To study if mobile elements in the *E. coli* K-12 and *Shigella flexneri* genome exhibit deviation in degrees of context dependency from the rest of the genome, we first implemented a controlled perturbation scheme for all predicted genes. Specifically, for each target position *i*, we masked that position and compared two scenarios: the original genomic context versus a context-swapped version. In the context-swapped condition, we replaced the local neighborhood (*w* = 5 genes on each side) surrounding the masked position with the corresponding neighborhood from a randomly selected distant position *j* (where |*i* −*j*| *>* 2*w* to ensure non-overlapping contexts). For each configuration, the model predicted the masked embedding, and we computed the mean squared error (MSE) between the predicted and ground truth embeddings as 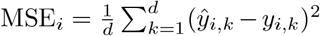, where *d* = 480 is the embedding dimension. The change in reconstruction error was quantified as 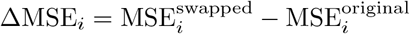, which serves as our context-dependency metric.

We extracted transposable element annotations for *Escherichia coli* and *Shigella flexneri* from NCBI GFF3 files. For each organism, we identified all genomic regions annotated with mobile element types, then extracted coding sequences (CDS) that overlapped these regions and whose product names contained “IS” (insertion sequence). This yielded organism-specific lists of IS-associated product terms used for subsequent filtering. All genes were ranked by their ΔMSE scores, and the bottom ones were designated as putative transposons. We used odds ratios (proportion of mobile elements in bottom 20% of all genes ranked by ΔMSE / proportion of mobile elements in rest of the genome) to quantify the likelihood that genes with low context-dependency scores correspond to true transposable elements.

### Genome scaffolding

To evaluate BacPT’s ability to reconstruct correct scaffold arrangements, each test genome’s ESM embeddings was partitioned into a specified number of scaffolds of approximately equal size. All possible permutations of scaffold order were generated, with each scaffold considered in both forward and reverse orientations. For each arrangement, protein embeddings were concatenated according to the scaffold order and padded to a maximum sequence length of 5000 positions. Random masking was applied to a specified proportion of positions in each embedding sequence with a masking fraction of 0.4. Masked positions were replaced with Gaussian noise sampled from a distribution parameterized by the mean and variance of the ESM embeddings of the training data. BacPT was used to reconstruct the masked embeddings for each arrangement. The MSE between predicted and true embeddings at masked positions was computed for each configuration. For computational efficiency, a random subset of up to 100 permutations per genome was evaluated when the total number of possible arrangements exceeded this threshold. To assess discriminative performance, native and permuted arrangements were treated as a binary classification problem, with native arrangements assigned positive labels and permuted arrangements negative labels. Reconstruction errors were negated to create scores where lower errors corresponded to higher values. Evaluations were performed across multiple scaffold counts (2, 3, 5, 7). A receiver operating characteristic curve (ROC) was generated by varying the classification threshold, and the area under the ROC curve was computed as the final performance metric, with results aggregated across all test genomes and scaffold configurations.

### Enzyme prediction pipeline

Enzyme activity profiles for 28 enzymes were curated from the BacDive database, with binary activity labels (positive/negative) derived from standardized API assays. Genomes were fetched from BacDive by filtering for strains with RefSeq accession numbers, yielding a total of 3,037 genomes. For each enzyme, its corresponding EC number was used to query CLEAN predictions across all genomes in the dataset, in order to determine if individual genomes harbor homologs of the enzyme-coding gene. Only strains for which a homolog was identified by CLEAN were retained for downstream analysis, restricting the prediction task to strains where the coding gene is present.

For each enzyme, embeddings of the focal enzyme-coding gene were extracted under three conditions: raw ESM embeddings, BacPT small-contextualized embeddings, and BacPT large-contextualized embeddings. For BacPT embeddings, the hidden state of the focal gene was extracted at each of the model’s layers independently, yielding a separate 480-dimensional representation per layer. A gene presence baseline was also computed, where precision and F1 score were derived directly from the fraction of homolog-containing strains exhibiting measurable enzyme activity, without any embedding-based model.

For each enzyme, embedding type, and layer, an *L*_2_-regularized logistic regression model was trained to predict binary enzyme activity. The dataset was split into 80% training and 20% test sets using stratified shuffle splitting to preserve class balance. The regularization strength *C* was selected via 3-fold cross-validation on the training set over a grid of values, optimizing for binary F1 score. The layer yielding the highest cross-validated F1 score on the training set was selected as the best layer, and the final model was retrained using that layer’s embeddings with the best *C*. Only enzymes with more than 10 positive and 10 negative examples were retained for evaluation. Performance was reported as the F1 score (*F* 1_original_) on the held-out test set, computed as:

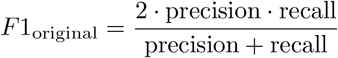

where precision and recall were computed from the confusion matrix on the test set. The entire procedure was repeated across five random seeds, and mean *F* 1_original_ was reported across seeds for each enzyme and embedding type. Gene synteny around the glutamate decarboxylase locus was visualized using the clinker tool [72].

### Operon classification task

Operon classification was performed using embeddings derived from BacPT and compared against baseline ESM embeddings. Operon annotations for *Escherichia coli* K-12 were obtained from RegulonDB [46], and operons containing at least three genes were selected for analysis.

To generate operon embeddings, we extracted representations from BacPT at the final layer for all genomic positions corresponding to genes within each operon. The embedding for an operon spanning positions from index *i*_start_ to *i*_end_ was computed as the mean across all positions: 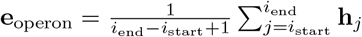, where **h**_*j*_ represents the 480-dimensional hidden state at position *j*. For ESM embeddings, the same averaging procedure was applied to the pre-computed protein-level representations. A balanced dataset was constructed by pairing each annotated operon with a randomly generated null operon of matched length, ensuring that null operons did not overlap any annotated operons by more than 50% and were sampled from valid genomic positions within the 4,297-protein *E. coli* K-12 genome.

Logistic regression with *L*_2_ regularization was employed to classify operons versus null regions. The dataset was split into 80% training and 20% testing sets with a fixed random seed for reproducibility. Hyperparameter tuning for the regularization strength *C* was performed via 3-fold cross-validation on the training set, and selecting the model with highest cross-validated accuracy. Model performance was evaluated on the held-out test set using F1 score and AUROC.

### Calculation of Jacobian matrices

To quantify pairwise gene interactions predicted by BacPT, we computed genome-wide interaction matrices using a perturbation-based approach adapted from the categorical Jacobian method [47]. For each protein position *i* in the genome, we masked the corresponding embedding by replacing it with a zero vector while keeping all other protein embeddings unchanged. The model was then used to generate predictions for all genomic positions under this perturbed input. The influence of the masked gene *i* on gene *j* was quantified as the *L*_2_ norm of the difference between the model’s prediction for position *j* with and without masking position *i*: 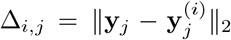, where y*_i_* represents the unperturbed prediction and 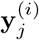 represents the prediction when position *i* is masked. This procedure was repeated for all *L* positions, yielding an *L*× *L* raw interaction matrix **M**.

To refine the interaction signal, we post-processed *M* using a series of transformations. First, each dimension was mean-centered independently by subtracting off the corresponding row and column means. We then took the absolute value of *M* entry-wise to enhance signal contrast: **M**← |**M**| . Symmetrization was achieved by computing **M**←**MM**^*T*^ to ensure that the interaction strength between genes *i* and *j* was consistent regardless of which gene was perturbed. Finally, we applied average product correction (APC) to remove background correlation biases: 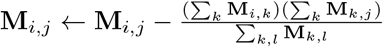 . Negative values resulting from APC were clipped to zero, and diagonal elements were set to zero to exclude self-interactions. We use the resulting processed matrix to represent the gene interaction predictions from BacPT.

### Correlation with STRING interactions

To validate the biological relevance of the predicted gene interactions, we compared the Jacobian matrix entries against interaction scores from the STRING database [48] for *E. coli* K-12. STRING provides probabilistic gene interaction scores derived from multiple evidence channels, including genomic neighborhood, gene fusion events, co-occurrence, text mining, coexpression, curated databases, and experimental evidence, as well as a combined score integrating all channels. For each evidence type, we extracted protein pairs along with their corresponding STRING scores and mapped them to positional indices in our genome ordering. To focus on high-confidence interactions and reduce the influence of spurious associations, we filtered protein pairs based on percentile thresholds of their STRING scores, testing thresholds at the 15th, 40th, 70th, and 90th percentiles. For each evidence type and threshold, we computed the Pearson correlation coefficient between the filtered STRING scores and the corresponding entries in our Jacobian matrix. The 90th percentile threshold was used for final reporting to ensure that only the most confident interactions from STRING were compared against our model’s predictions.

### Gene cluster identification

Gene cluster annotations were retrieved from the Minimum Information about a Biosynthetic Gene Cluster (MiBiG) database [49]. To ensure the robustness of the functional signals, we filtered the dataset to include only BGCs containing ≥5 genes. Our analysis focused on the top eight taxa with the highest number of BGCs per genome, resulting in a finalized set of 82 BGCs for evaluation. For each BGC, we calculated an enrichment ratio (*R*) using the Jacobian matrix generated from BacPT:

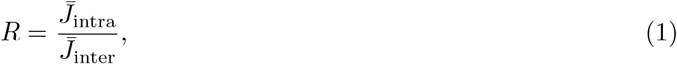

where 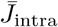 represents the median Jacobian score among genes within the cluster, and 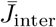 represents the median score between genes inside the cluster and those in the surrounding genomic background.

To assess the statistical significance of these ratios, we generated a null distribution for each BGC by randomly sampling contiguous genomic windows with the same number of genes from the same genome as the test BGC. Each BGC was then assigned a percentile rank relative to its specific null distribution. A high percentile rank indicates that the constituent genes of a BGC exhibit higher coupling with each other than with genes outside the cluster, suggesting a specialized evolutionary and functional modularity captured by BacPT.

### Trait prediction pipeline

Phenotypic data for 3,037 bacterial genomes were retrieved from the BacDive database by filtering for strains with RefSeq accession numbers. All available binary cellular phenotypes were included except enzymatic activities, which were analyzed separately. Categorical labels were mapped to binary values, with ambiguous or unspecified values discarded. Traits were retained only if both positive and negative classes contained at least 10 samples in both training and test sets, yielding a final dataset of 66 traits spanning metabolite utilization, biosynthesis, antibiotic resistance, and growth conditions. The dataset was split into 80% training and 20% test sets using stratified shuffling to preserve class balance.

For each genome, mean BacPT embeddings were computed by averaging the hidden states across all protein positions at each layer independently, yielding a 480-dimensional representation per layer per genome. Mean ESM embeddings were computed analogously by averaging raw ESM embeddings across all proteins in the genome. A Pfam presence/absence feature vector was constructed for each genome by running InterProScan against all predicted proteins using the Pfam database, and encoding the result as a binary vector of length 8,552 indicating the presence or absence of each Pfam domain.

For each trait and embedding type (ESM, BacPT-large, BacPT-small, Pfam presence/absence), an *L*_2_-regularized logistic regression model was trained using grid search with 5-fold cross-validation optimizing average precision. For BacPT embeddings, models were trained independently for each layer, and the layer yielding the highest F1 score on the training set was selected as the best layer for final evaluation. Performance on the held-out test set was reported as F1 score. Traitar predictions were obtained by running the Traitar pipeline on the same set of genomes, and its predictions were compared against ground truth labels on the intersecting set of 17 traits available in both BacDive and Traitar.

### Trait attribution analysis

To identify which protein domains (Pfam families) contribute most to phenotype predictions, we performed a perturbation-based attribution analysis on the trained BacPT models by ablating proteins containing each Pfam domain and measuring the resulting change in model predictions. For each genome, we computed a baseline prediction by forward-passing the complete set of ESM embeddings through the BacPT models, extracting the mean-pooled representation from the best-performing layer (determined via cross-validation during training), and obtaining the raw logit from the trained logistic regression classifier.

To assess each Pfam domain’s contribution, we replaced the ESM embeddings of all proteins containing that domain with zero vectors (which effectively removes their information content since embeddings were normalized to mean zero and standard deviation one), computed the perturbed logit, and calculated attribution metric: the total effect Δ_total_ = logit_baseline_ −logit_perturbed_. Positive Δ values indicate that removing the domain decreases the probability of the positive class.

We aggregated results across genomes by computing, for each Pfam domain, the mean and standard error of Δ_total_. The analysis was implemented in PyTorch with batch processing (batch size = 32), mixed-precision inference (float16), and focused on genomes with positive phenotype labels to identify domains supporting positive predictions, with Pfam-to-protein mappings obtained from InterProScan annotations.

### Ecological interaction prediction pipeline

To evaluate BacPT’s ability to capture ecological interactions between microbial species, we predicted interaction types from pairwise genome embeddings using a logistic regression framework. Interaction data were obtained from [55], which catalogs pairwise microbial interactions across various carbon source environments. For each species pair, we extracted genome-wide embeddings from BacPT at each layer by computing the mean embedding across all protein positions in the genome: 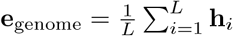, where *L* is the number of proteins and **h**_*i*_ represents the 480-dimensional hidden state at position *i*. For a given species pair (*A, B*), we constructed a feature vector by concatenating the genome embeddings of both species and their element-wise product: **x** = [**e**_*A*_; **e**_*B*_; **e**_*A*_ ⊙ **e**_*B*_]. The element-wise product was included to capture potential synergistic features between the two genomes. The feature vectors for ESM-based models were constructed in the same manner. Duplicate species pairs within the same environment were removed to prevent data leakage.

We trained separate *L*_2_-regularized logistic regression models for each carbon source environment to classify interaction types (e.g., mutualism, competition, parasitism) from these feature vectors. To ensure robustness, we filtered out interaction classes with fewer than three instances per environment. The regularization strength *C* was selected via grid search using stratified *k*-fold cross-validation with *k* = 3 on the training set, optimizing for macro-averaged F1 score. Two distinct data splitting strategies were employed to assess model generalization. In the *random split* strategy, species pairs were randomly divided into training and validation sets (80/20 split), allowing the same species to appear in both sets but ensuring that specific pairs were unique to each set. Stratified k-fold cross-validation (*k* = 3) was performed on the training set for hyperparameter tuning, and performance was evaluated on the held-out validation set. In the *species-disjoint split* strategy, species themselves were randomly partitioned into training and validation sets (80/20 split), ensuring complete separation such that no species in the validation set appeared during training. This more stringent evaluation tested the model’s ability to generalize to entirely novel species. For each split, we verified that all interaction classes were represented in both training and validation sets; if this condition was not met after 300 random shuffles, that split was discarded. Due to potential class imbalance in the training set after species partitioning, we adaptively selected the number of inner cross-validation folds based on the minimum class count, using 2-fold CV when the smallest class had fewer than three samples.

Both splitting strategies were repeated across five random seeds to account for variability in data partitioning. For each seed, model type, layer, and environment, we computed the macro-averaged F1 score. The final reported performance for each configuration was the mean F1 score across all five seeds, providing a robust estimate of the model’s predictive capability for microbial ecological interactions.

## Code availability

BacPT model weights and code to reproduce the analyses are available at: https://github.com/palsetuf/BacPT.

## Acknowledgement

Research reported in this publication was supported by the National Institute of General Medical Sciences of the National Institutes of Health under award number R35GM154908, and University of Florida College of Liberal Arts & Sciences.

## 1 Supplementary information

### 1.1 Supplementary methods

### Supplementary Figures

**xFigure S1.**
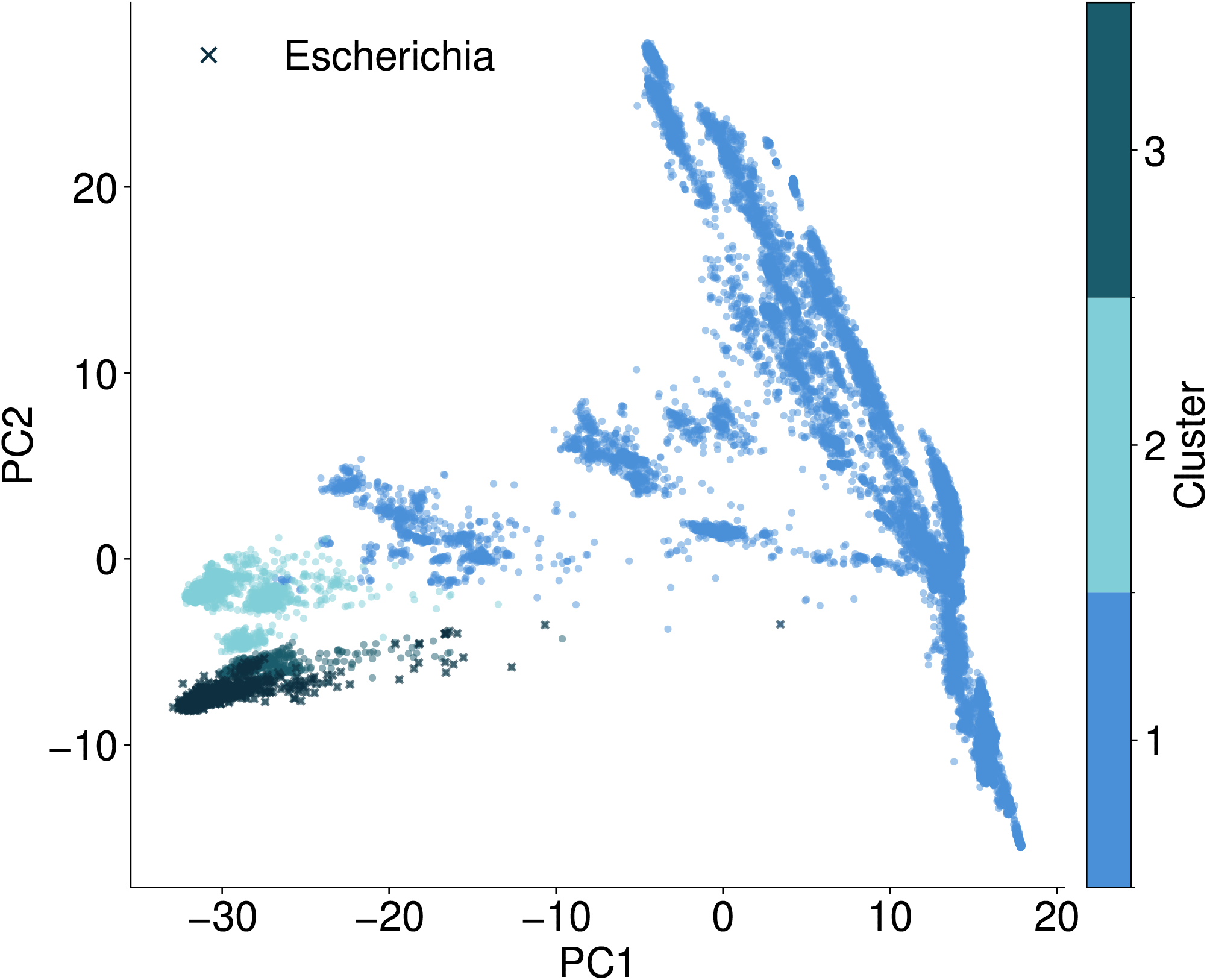
BacPT training data was selected based on clustering analysis of protein family representations of 33,140 bacterial genomes. The cluster 3, including the genus Escherichia (teal) was chosen as the test set. Final training data consists of 28,133 genomes.

**Figure S2.**
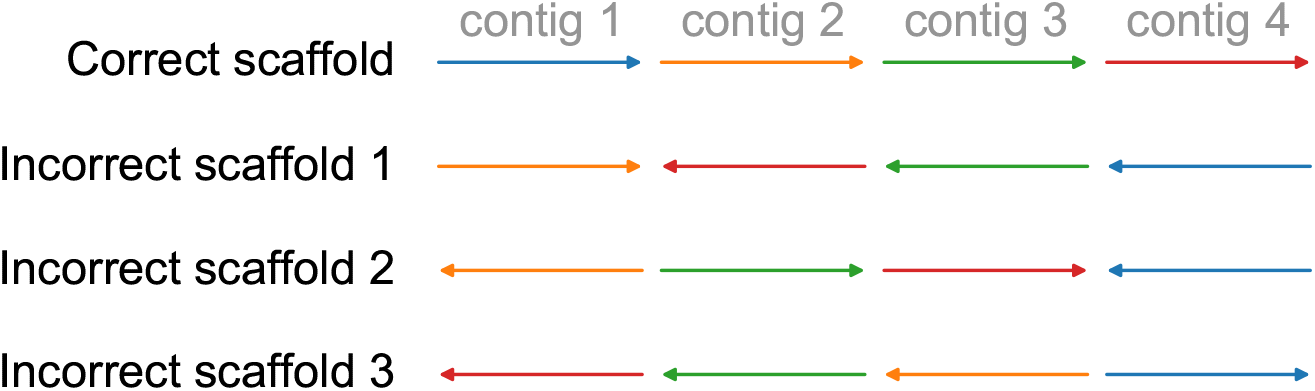
Simulated genome scaffolding task. A test bacterial genome is segmented into 2-7 nonoverlapping contigs of roughly equal sizes. Incorrect scaffolds were generated by randomly permuting the orientation and the order of the contigs. BacPT masked reconstruction MSE was then used to differentiate correct vs. incorrect scaffolds.

**Figure S3.**
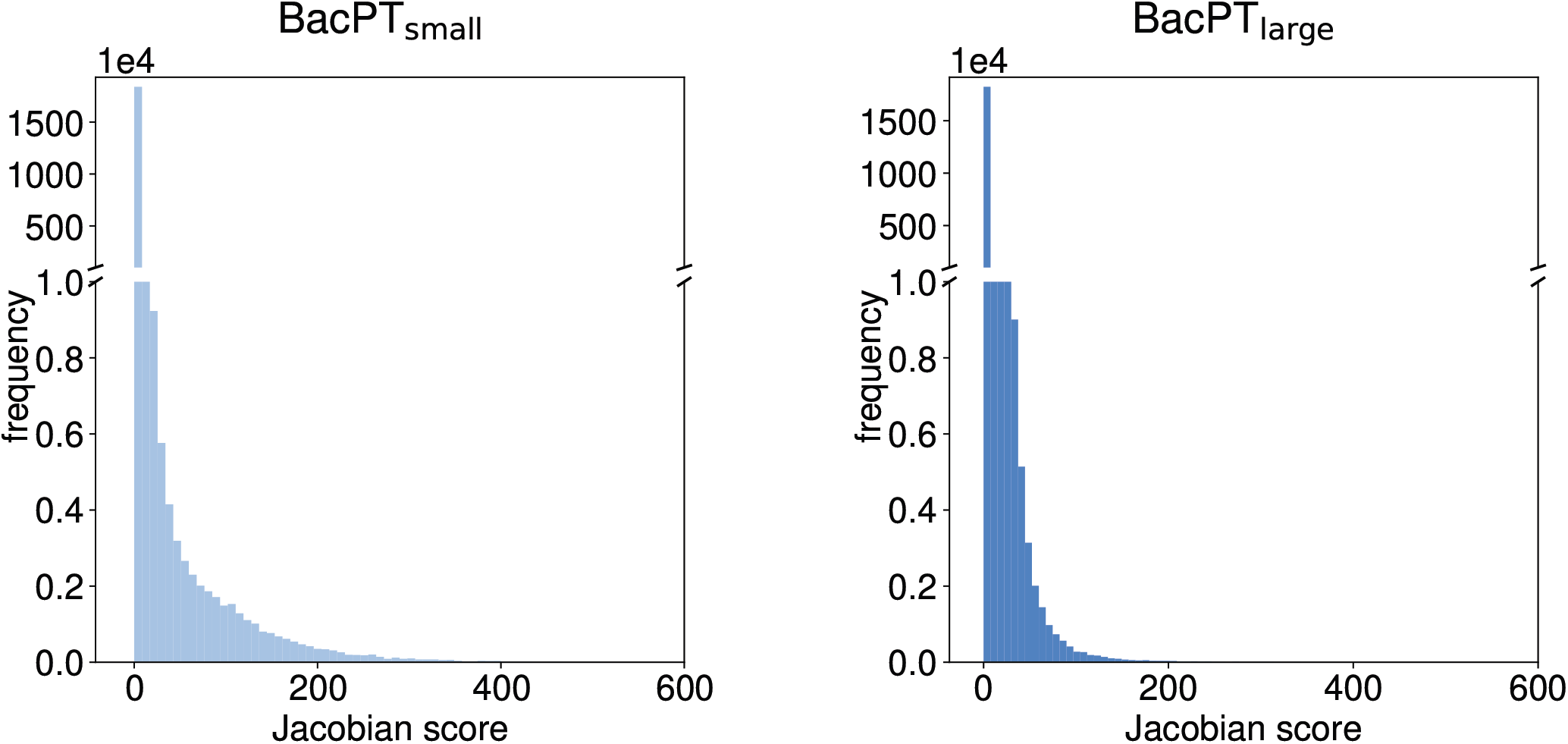
Distribution of interaction strengths between genes in the symmetrized Jacobian matrix *M* .

**Figure S4.**
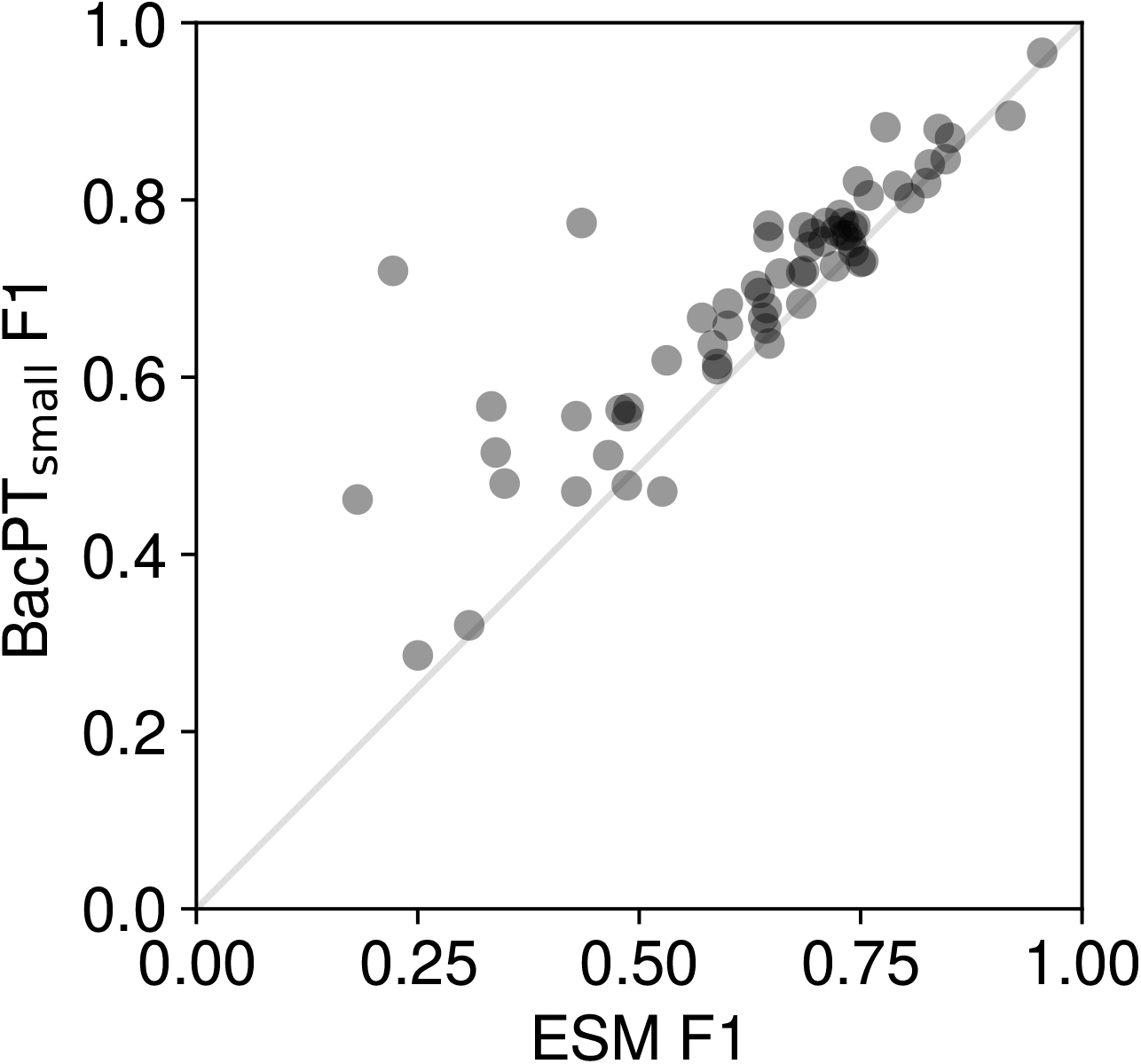
Comparison of F1 scores on trait prediction tasks between linear probes trained using BacPT-small embeddings and ESM embeddings.

**Figure S5.**
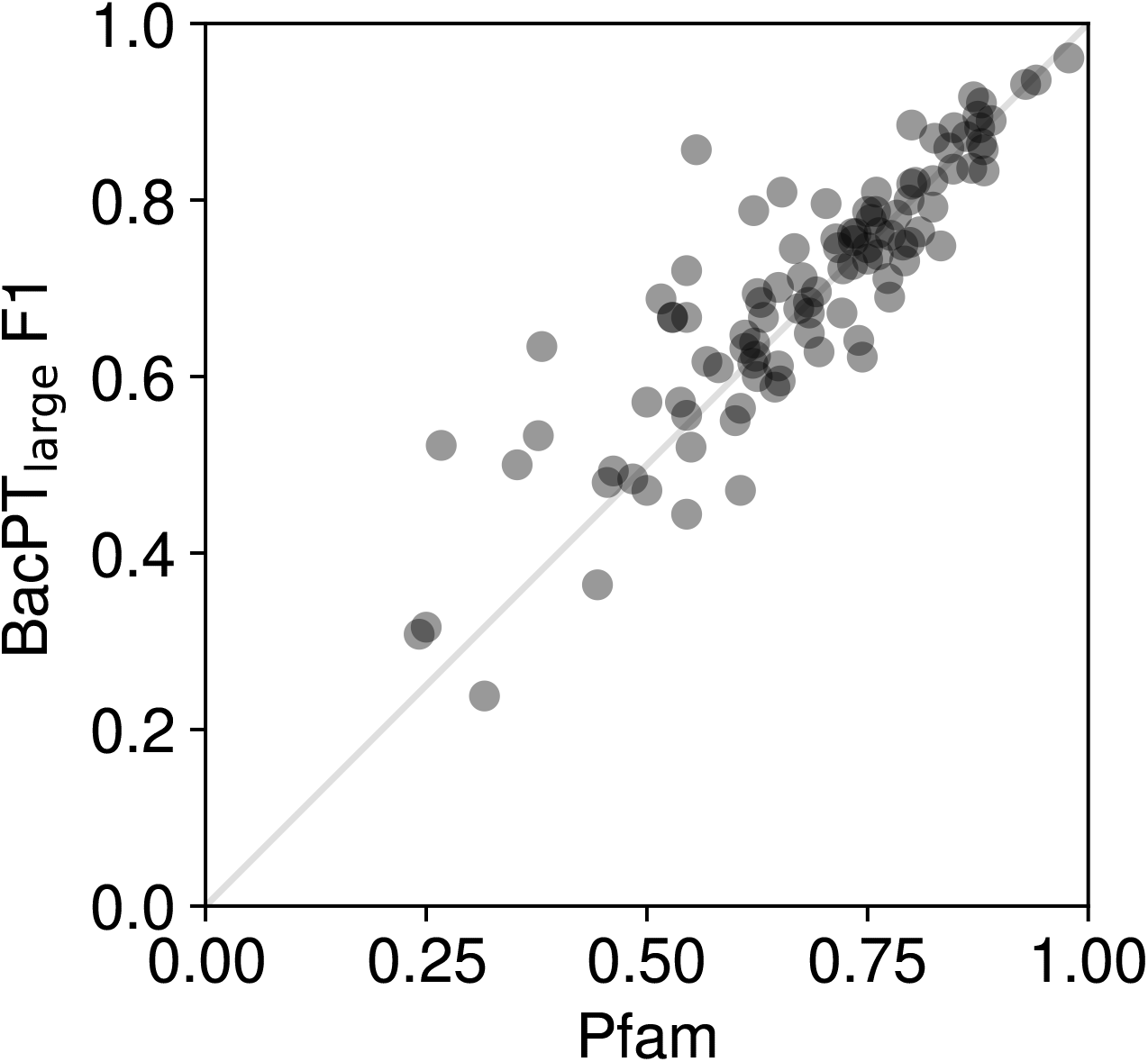
Comparison of F1 scores on trait prediction tasks between linear probes trained using BacPT-large embeddings and models trained using Pfam presence/absence.

